# Electric-field induced sleep promotion and lifespan extension in Gaucher’s disease model flies: Association with improving ER stress and autophagy

**DOI:** 10.1101/2024.04.01.587515

**Authors:** Takaki Nedachi, Haruhisa Kawasaki, Eiji Inoue, Takahiro Suzuki, Yuzo Nakagawa-Yagi, Norio Ishida

**Affiliations:** Hakuju Institute for Health Science Co., Ltd., 1-37-5 Tomigaya, Shibuya-ku, Tokyo 151-0063, Japan; Institute for Chronobiology, Foundation for Advancement of International Science (FAIS), 3-24-16 Kasuga, Tsukuba, Ibaraki 305-0812, Japan; Tokyo Research Center, Kyushin Pharmaceutical Co., Ltd., 1-22-10 Wada, Suginami-ku, Tokyo 166-0012, Japan; SHIGRAY Inc., 14-4-A2 Kitaarakawaoki, Tsuchiura, Ibaraki 300-0876, Japan; Tokyo Kasei University, 1-18-1 Kaga, Itabashi-ku, Tokyo 173-8602, Japan

**Keywords:** mitophagy, neurodegeneration, Parkinson’s disease (PD), extremely low frequency (ELF)

## Abstract

Gaucher’s disease (GD), one kind of genetic disease, is characterized by a mutation in metabolic enzyme, glucocerebrosidase (GBA), leading to accumulation of its substrate, glucosylceramide in tissues. We previously discovered that the *minos*-inserted mutation of the *GBA* gene in fruit flies, *Drosophila melanogaster*, reproduced human GD characteristics and can be used as a promising model for molecular mechanisms study. Recently, we reported that extremely low-frequency electric fields (ELF-EFs) promoted sleep and extended the lifespan of normal flies.

In this study, GD model flies were exposed to ELF-EFs, and sleep parameters, gene expressions, and longevity were evaluated. The amount of sleep and lifespan increased after EF exposure. Simultaneously, the expression of the endoplasmic reticulum (ER) stress-related gene, *PERK* and the autophagy-related gene, *p62* was elevated. The data suggest that the healthy effects of EFs on GD flies are associated with improved adaptation to cellular stress.

## Introduction

Gaucher’s disease(GD) is caused by mutations in the glucocerebrosidase (GBA) gene, decreased enzymatic activity of GBA, and accumulation of its substrate, glucosylceramide, in cells and tissues, leading to a variety of symptoms in the bone, blood, internal organs, and nervous system (Chen and Wang, 2008; Nagral, 2014). Mutants of the GBA gene are known to increase the risk of Parkinson’s disease (PD), suggesting a common molecular mechanism(s) between GD and PD (Alcalay et al., 2014; Mitsui et al., 2009). Owing to the abundance of genetic tools and ease of housing, pathological model animals with human GD symptoms have been established in GBA mutant fruit flies (Kawasaki et al., 2017; Kinghorn et al., 2016; Suzuki et al., 2013). We previously reported that the expression of the human mutated glucocerebrosidase gene (*hGBA*), which is associated with neuronopathy in GD patients, causes neurodevelopmental defects in *Drosophila* eyes. We showed that endoplasmic reticulum (ER) stress is elevated in *Drosophila* eyes carrying mutated hGBA by using of the ER stress markers dXBP1 and dBiP. We also found that ambroxol, a potential pharmacological chaperone for mutated hGBA, could alleviate the neuronopathic phenotype by reducing ER stress. This GD model is accompanied by elevated ER stress, unfolded protein response (UPR), and inflammation, which are the main therapeutic targets of drugs (Cabasso et al., 2019; Kawasaki et al., 2017; Kinghorn et al., 2016; Suzuki et al., 2013). In our previous report, a *minos* insertion mutant with a homologous GD gene (*CG31414*) accumulated hydroxy-glucosylceramide in the whole body of *Drosophila melanogaster*. This *minos* mutant showed abnormal phenotypes of climbing ability and sleep, and a short lifespan. These abnormal phenotypes are similar to those of GD in humans (Cabasso et al., 2019; Kawasaki et al., 2017).

Compared with surgical or pharmaceutical approaches (Nagral, 2014; Sam et al., 2021), physical therapy for GD is not known. Recently, an extremely low-frequency electric field (ELF-EF) was reported to promote sleep and extend the lifespan of normal flies (Kawasaki et al., 2021), suggesting that a physical approach may be an alternative to medication. In this study, *minos* mutant GD model flies were exposed to ELF-EFs, and sleep parameters, longevity, and related gene expression were evaluated.

## 2. Materials and methods

### 2.1 Fly strains

The fruit fly *Drosophila melanogaster* was used in the experiments. Flies with a *CG31414* gene mutation, which was established as a *GBA* mutant, were used as the GD model (Kawasaki et al., 2017). w^1118^, the parental line of the GD mutant, was used as a healthy control. Flies were reared with standard cornmeal food in a temperature-controlled incubator at 25 °C under 12:12 h light/dark (LD) cycles.

### 2.2 EF exposure system

The same exposure system was used in a previous study (Kawasaki et al., 2021). Briefly, parallel plate electrodes of 300 mm × 300 mm stainless steel were pasted on the polyvinyl chloride plates; one was applied with a 50 Hz alternating current (AC) high voltage and the other was connected to the ground (0 V). Four Teflon pillars with a radius of 20 mm were used as spacers between parallel electrodes.

A high voltage device, Healthtron HEF-P3500 (Hakuju Institute for Health Science, Tokyo, Japan), was used for voltage application (Kawasaki et al., 2021).

EF is defined as the electric voltage per length (V/m) and is dependent on the voltage and length between the electrodes. In this experiment, an output voltage of 3.5 kV and pillar length of 100 mm were employed; thus, a 35 kV/m EF was generated.

An identical pair of parallel electrodes was set for sham exposure, where both electrodes were connected and grounded to ensure a zero EF.

### 2.3 Sleep monitoring

The *Drosophila* Activity Monitoring System (TriKinetics, Waltham, MA, USA) was used for sleep monitoring (Kawasaki et al., 2017; Kawasaki et al., 2021). The activity count was measured using a 1-min bin, and an inactive period ≥ 5 min was defined as sleep.

Each fly around 7-day-old was placed in a glass tube of 2 mm diameter, 50 mm length with one end filled with 5 % sucrose, 2% agar medium (Fig. 1a). After resting for one night, glass tubes containing the flies were placed inside the electrodes, and the EF was exposed for 9 h during the daytime (Figs. 1a and 1b). The tubes were collected immediately after EF exposure, and the subsequent nighttime sleep was monitored for 12 h (Fig. 1b) in the incubator. Sleep bout number, total sleep, and sleep episode duration were evaluated.

**Fig. 1.**
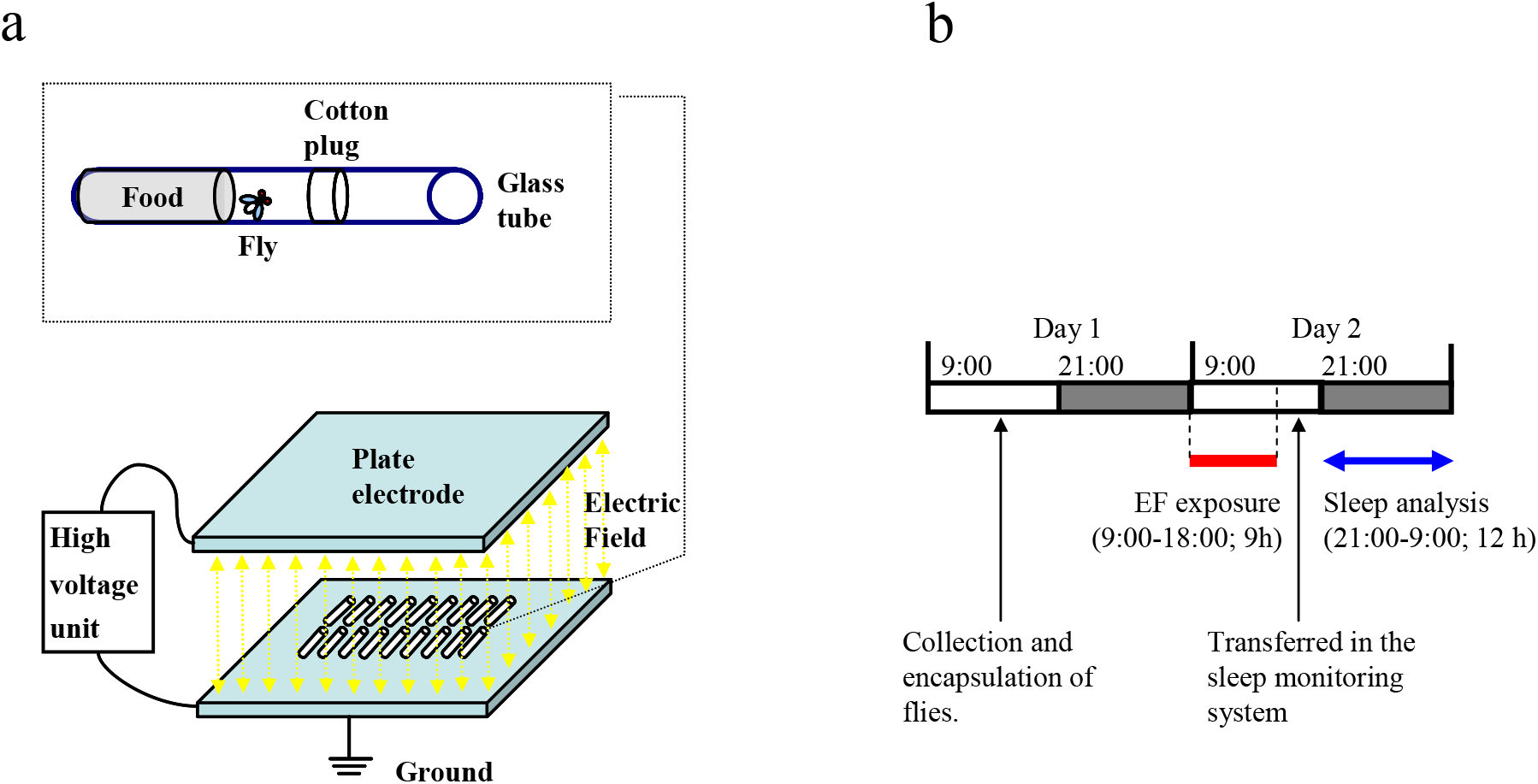
Electric field (EF) exposure system and sleep measurement schedule. (a) An EF was generated by applying a high voltage (3.5 kV, 50 Hz) between the two parallel plate electrodes (300 mm × 300 mm) separated by a distance of 100 mm. (b) Flies were collected and put into the tubes during daytime on day 1. EF exposure was conducted for 8 h; from 9:00 to 18:00 during the daytime on day 2. The tubes were then transferred to the sleep monitoring device to measure the following nighttime sleep.

### 2.4 Lifespan assay

Vials with low-nutrition medium containing 5 % glucose and 1 % agar were prepared. Newly eclosed male flies were collected under anesthesia within 48 h after eclosion and transferred into vials for lifespan assays (Kawasaki et al., 2017; Kawasaki et al., 2021). Flies’ longevity was measured under continuous EF/Sham exposure conditions in a temperature-controlled room at 25 °C under 12:12 h LD cycles. The medium was changed at least twice a week. Each vial contained 20 flies, and 79-80 flies were used for each group.

### 2.5 qRT-PCR

Flies treated with the same procedure as the lifespan assay were collected after 7-day EF/Sham exposure and homogenized in RNAiso reagent (Takara Bio, Shiga, Japan). Ten to twenty flies were homogenized in one tube, and 4 – 8 biological replicates were prepared for one group. RNA was extracted, and qRT-PCR was performed as previously described (Kawasaki et al., 2017). An autophagy-related gene *p62*, ER-related genes *BiP, Ire1, PERK, CG14715*, a PD-related gene *PINK1*, a lysosome-related gene *cathD* (*cathepsin D*), and a bone morphogenetic protein (BMP) signaling-related gene *tok* were evaluated. *RPL32* (*ribosomal protein L32*) gene was used as an internal control. Primer sequences are listed in Supplementary Table 1.

### 2.6 Statistical analysis

The significance of differences in sleep and qRT-PCR data was estimated using Student’s t-test. Longevity was evaluated using the log-rank test. Differences were considered statistically significant at P < 0.05.

## 3. Results

### 3.1 Changes in sleep status in GD flies by EF exposure

Sleep bout number in EF group was significantly decreased, while total sleep and sleep episode duration were significantly increased in GD flies (Fig. 2b). In contrast, these sleep parameters were not significantly changed in the control, w^1118^ flies (Fig. 2a).

**Fig. 2.**
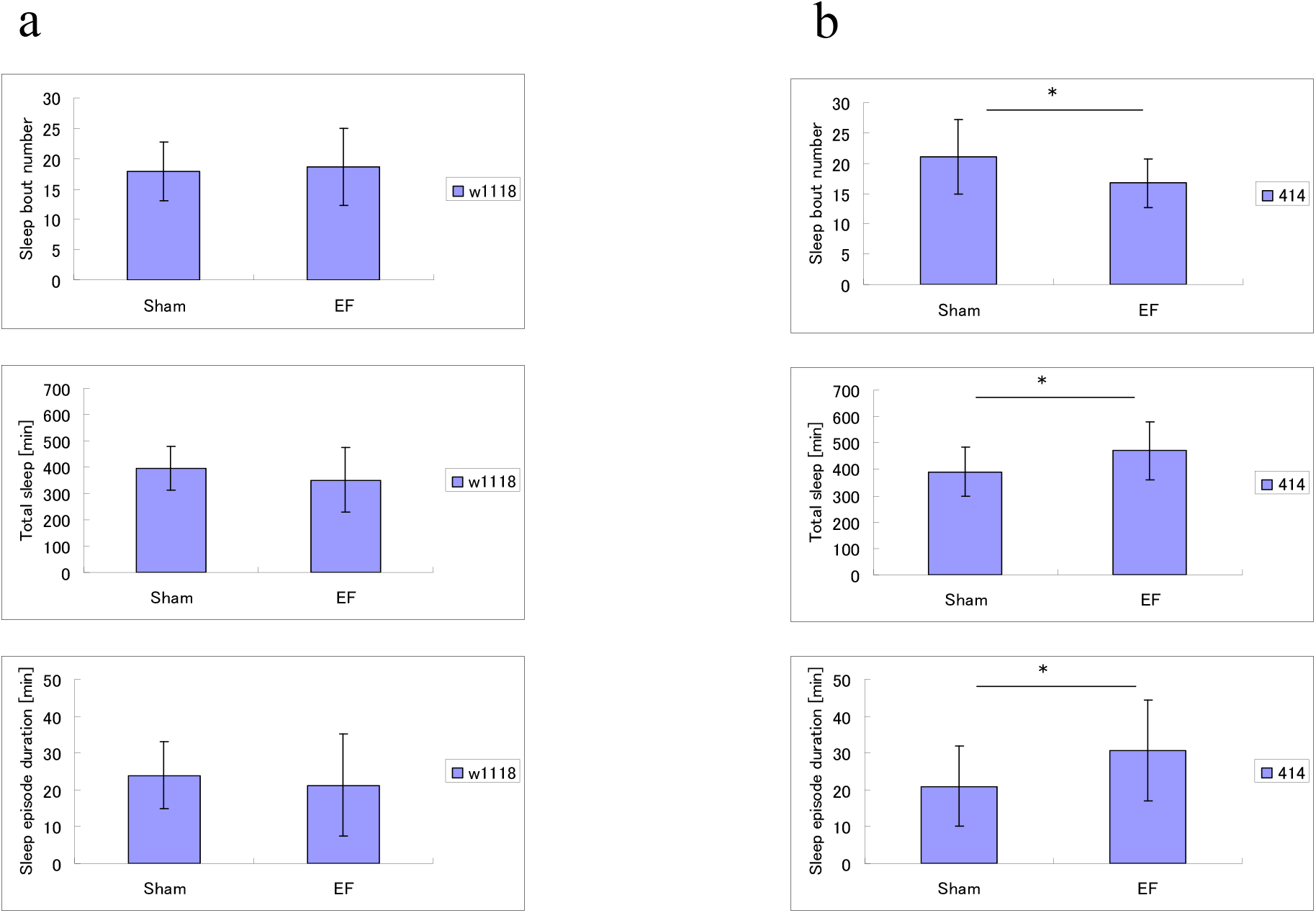
Effects of the EF exposure on the sleep parameters in healthy control (w^1118^) and GD flies. In w^1118^ flies (a), sleep bout number (top), total sleep (middle), and sleep episodes duration (bottom) during nighttime were not changed by the EF exposure. In GD flies (b), sleep bout number was decreased, while total sleep time and sleep episode duration were significantly increased in the EF group. Statistical data are expressed as the mean ± S.D. * p < 0.05 by Student’s t-test. N = 31 flies each for w^1118^; n = 14 - 18 flies each for GD.

### 3.2 Effects of EF exposure on the lifespan of GD flies

The median survival time of GD flies in EF and Sham groups was 11 days, whereas it was 22 (EF group) - 27 (Sham group) days in healthy flies (Figs. 3a and 3b).

**Fig. 3.**
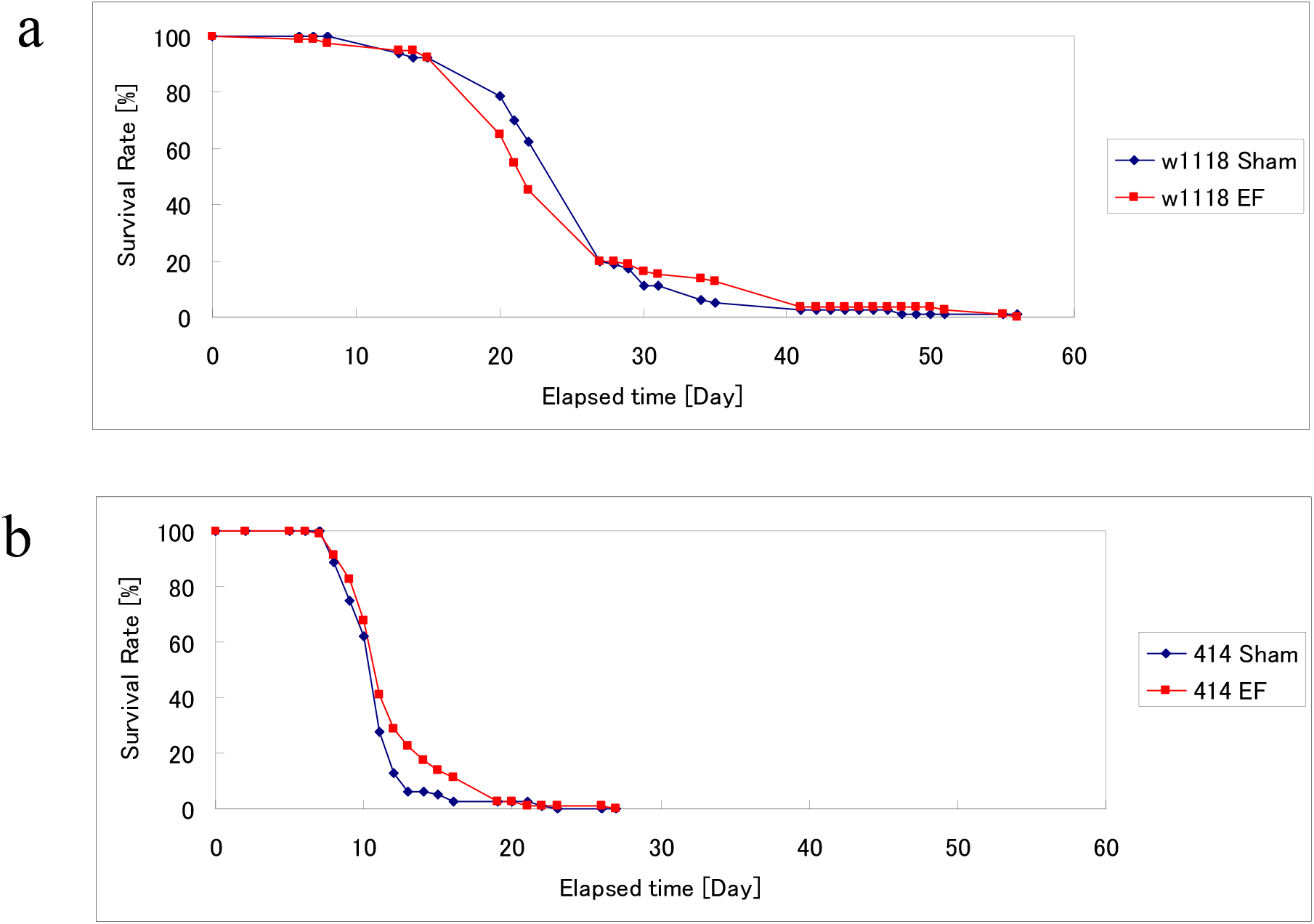
Effects of the EF exposure on longevity. (a) The lifespan of w^1118^ flies (n = 80) did not change after the EF exposure (p = 0.548). (b) The lifespan of GD flies (n = 79-80) was half that of the healthy flies (n = 80). EFs significantly extended the lifespan in GD flies (p = 0.029). Statistical analyses were performed using the log-rank test.

The lifespan of GD flies was significantly longer in EF group than in Sham group (p = 0.029 (Fig. 3b)). In contrast, the lifespan of healthy flies was not significantly changed (p = 0.548 (Fig. 3a)).

### 3.3 Changes in gene expression in GD flies by EF exposure

Upregulation in autophagy marker *p62* was observed in GD flies in comparison with the normal ones (Fig. 4a). The other genes such as ER stress marker, *PERK* was not significantly altered (Fig. 4b). *p62* and *PERK* were significantly upregulated in GD flies after the 7-day EF exposure (Figs. 5a and 5b). In contrast, the expression of the other genes (*BiP, Ire1, CG14715, PINK1, cathD*, and *tok*) was not significantly changed in EF vs. Sham group (Supplementary Fig. 1).

**Fig. 4.**
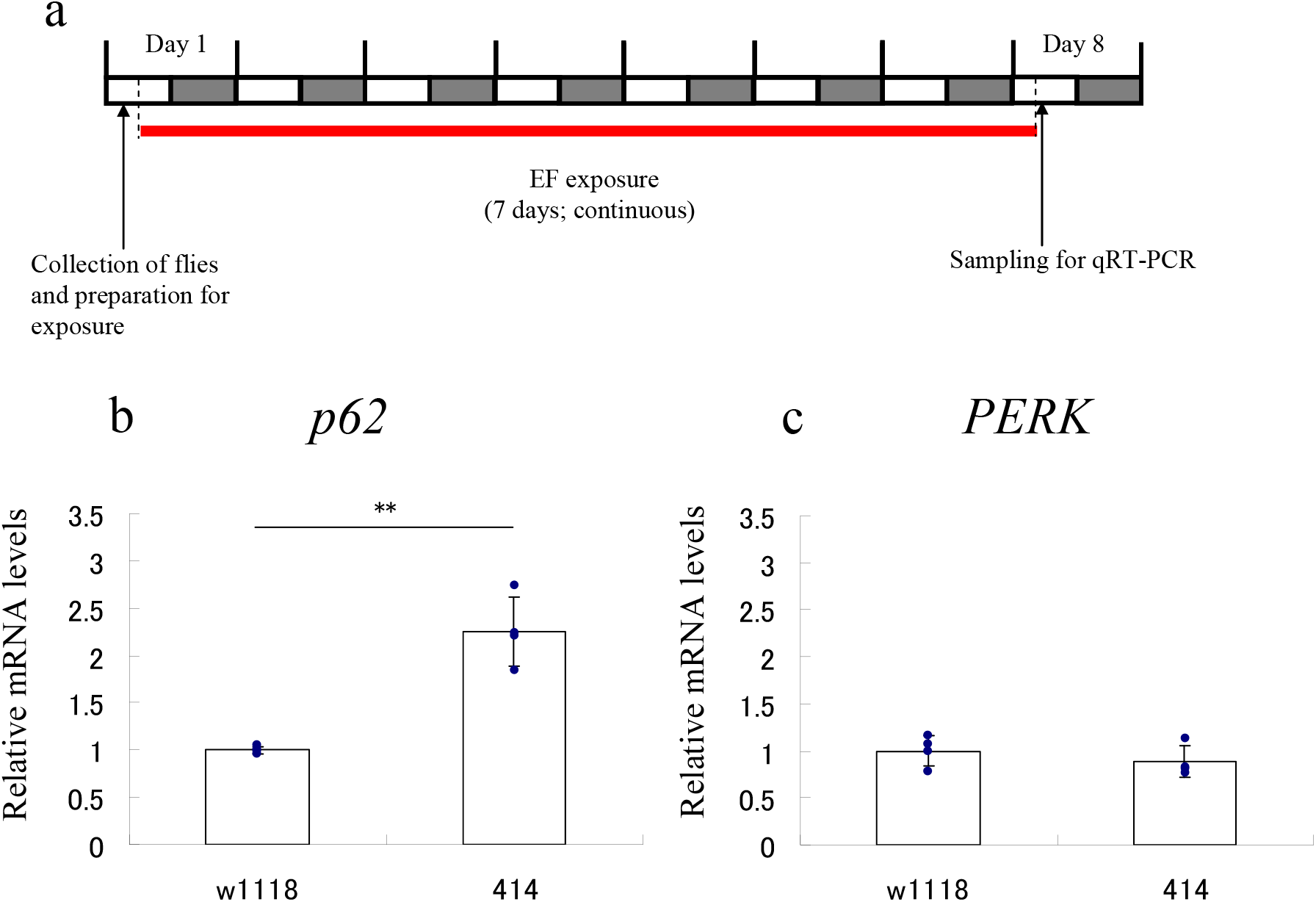
Gene expression assay using qRT-PCR. (a) The EF exposure was conducted continuously for 7 days. (b) Expression of the autophagy-related gene *p62* was significantly higher in GD flies than in healthy flies. (c) In contrast, other genes such as *PERK* were not altered. Statistical data are expressed as the mean ± S.D. Statistical analysis was conducted using Student’s t-test. * p < 0.05; ** p < 0.01. N = 4 biological replicates per group.

**Fig. 5.**
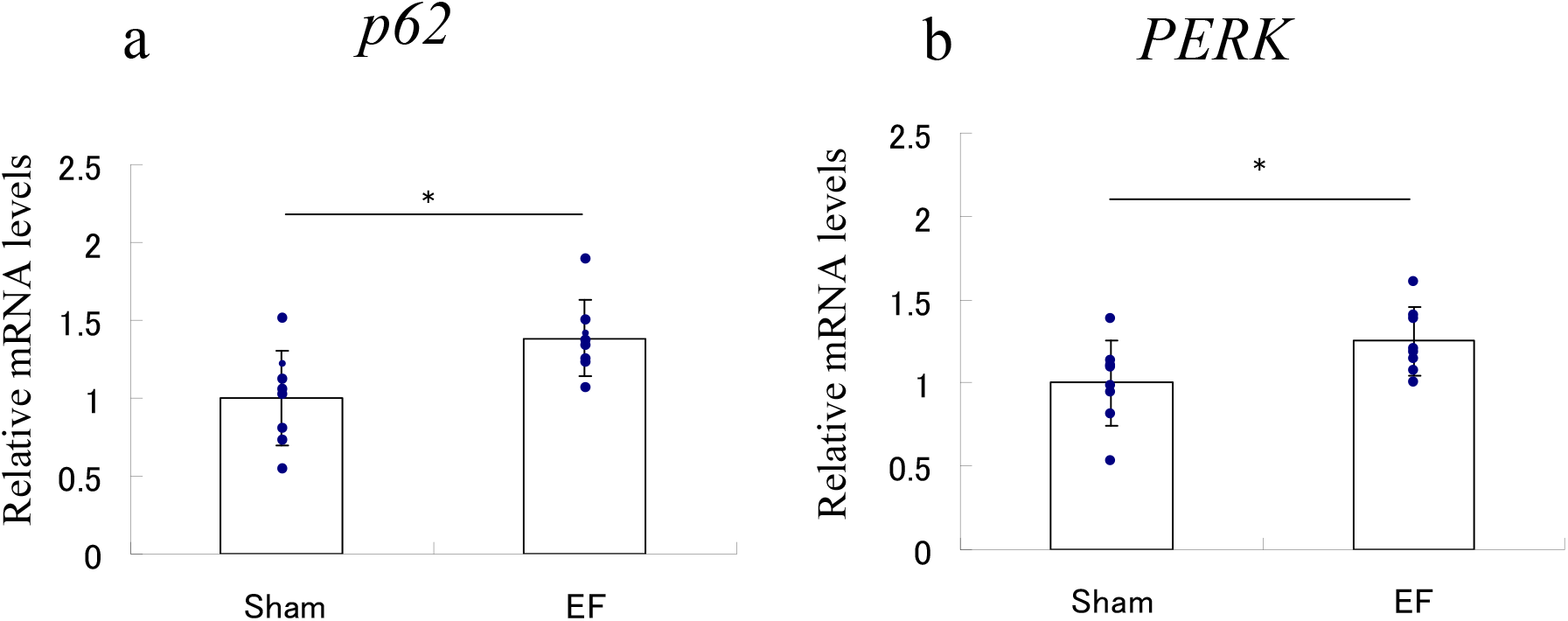
Altered gene expression by the EF exposure in GD flies. The 7-day EF exposure significantly elevated the expression of *p62* (a) and *PERK* (b). Statistical data are expressed as the mean ± S.D. Statistical analysis was conducted using Student’s t-test. * p < 0.05; ** p < 0.01. N = 8 biological replicates per group.

## 4. Discussion

This study showed that EFs affect sleep improvement, lifespan extension, and changes in the gene expression in GD model flies.

Among the tested genes, *PERK* and *p62* levels were increased by EF exposure. *p62* upregulation in GD vs. normal flies was also reproduced in this study (Kawasaki et al., 2017). Furthermore, EFs extended longevity and ameliorated GD, which was associated with increased *p62* expression. These data suggest that *p62* upregulation prolongs lifespan by improving mitochondrial function and mitophagy (Aparicio et al., 2019). In addition to autophagic/mitophagic process, *p62* is known to activate the Kelch-like ECH-associated protein 1 (Keap1) - Nuclear factor erythroid 2-related factor 2 (Nrf2) pathway (Ichimura et al., 2013) and upregulate numerous cytoprotective genes for cell survival. Therefore, EF-induced *p62* upregulation in GD flies may be an adaptation to cellular stress for survival in GD flies.

The interaction between *p62* and sleep is still controversial, as sleep deprivation either increased (Cao et al., 2021) or decreased (Li et al., 2020; Yang et al., 2019) p62 protein levels in several tissues.

PERK may activate the p62-mediated non-canonical KEAP1-Nrf2 pathway and may play a protective role against lipotoxic stress (Lee et al., 2022). Therefore, the co-upregulation of *p62* and *PERK* observed in our study may decrease ER stress in lipid metabolism. Furthermore, PERK activation by ER stress has been shown to promote sleep in nematodes (Kawano et al., 2023). PERK overexpression in flies induced sleep, where wake-promoting neuropeptide expression was suppressed (Ly et al., 2020). Thus, EFs may increase sleep, at least in part, by activating the PERK pathway.

As shown in our previous report on wild-type flies, Oregon R (Kawasaki et al., 2021), EF exposure had healthy effects on sleep and longevity in GD flies. However, these parameters were not significantly changed in normal control, w^1118^ flies. Recently, a shorter lifespan and weakness in response to various stressors have been reported in w^1118^ flies (Ferreiro et al., 2018). Thus, w^1118^ flies may differ from Oregon R flies, leading to unchanged sleep and lifespan following EF exposure.

GBA gene deficiency promotes protein aggregation and is linked to neurodegenerative diseases such as PD (Aflaki et al., 2017; Davis et al., 2016; Do et al., 2019; Maor et al., 2016; Vieira et al., 2022). Recently, EFs have been shown to prevent abnormal protein aggregation in solution (Pandey et al., 2019). If this occurs *in vivo*, EFs may alleviate GD symptoms by preventing protein aggregation. Our previous study showed that EF treatment elevated N-palmitoyl serine levels in healthy humans, which extended the lifespan of PD model flies (Nakagawa-Yagi et al., 2021). Further studies using *Drosophila* models of neurodegenerative diseases, such as PD and GD, should be conducted for clarifying the effect of ELF-EFs.

Further evidence is needed to explain why ELF-EFs induce sleep promotion and lifespan extension in GD model flies at a molecular level.

## Abbreviations

ELF: extremely low frequency
EF: electric field
GD: Gaucher’s disease
PD: Parkinson’s disease
GBA: glucocerebrosidase
ER: endoplasmic reticulum
UPR: unfolded protein response
XBP1: X-box binding protein 1
BiP: binding immunoglobulin protein
LD: light/dark
PERK: protein kinase R-like
ER: kinase
Ire1: inositol-requiring enzyme 1
PINK1: PTEN-induced kinase 1
cathD: cathepsin
D: RPL32, ribosomal protein
L32: Keap1, Kelch-like
ECH: associated protein 1
Nrf2: NF-E2-related factor 2

## CRediT authorship contribution statement

Takaki Nedachi: Conceptualization, Methodology, Formal analysis, Investigation, Writing – original draft. Haruhisa Kawasaki: Conceptualization, Methodology, Formal analysis, Investigation.

Eiji Inoue: Conceptualization, Methodology, Formal analysis, Investigation. Takahiro Suzuki: Conceptualization, Investigation. Yuzo Nakagawa-Yagi: Conceptualization, Investigation. Norio Ishida: Supervision, Conceptualization, Methodology, Writing & editing, Project administration.

## Data availability

The data that has been used is confidential.

## Acknowledgments

We thank Mr. Akikuni Hara (Hakuju Institute for Health Science Co., Ltd., Tokyo, Japan) for his advice and help.

