## Supplementary data for "Electric-field induced sleep promotion and lifespan extension in Gaucher’s disease model flies: Association with improving ER stress and autophagy"

### Slide 1
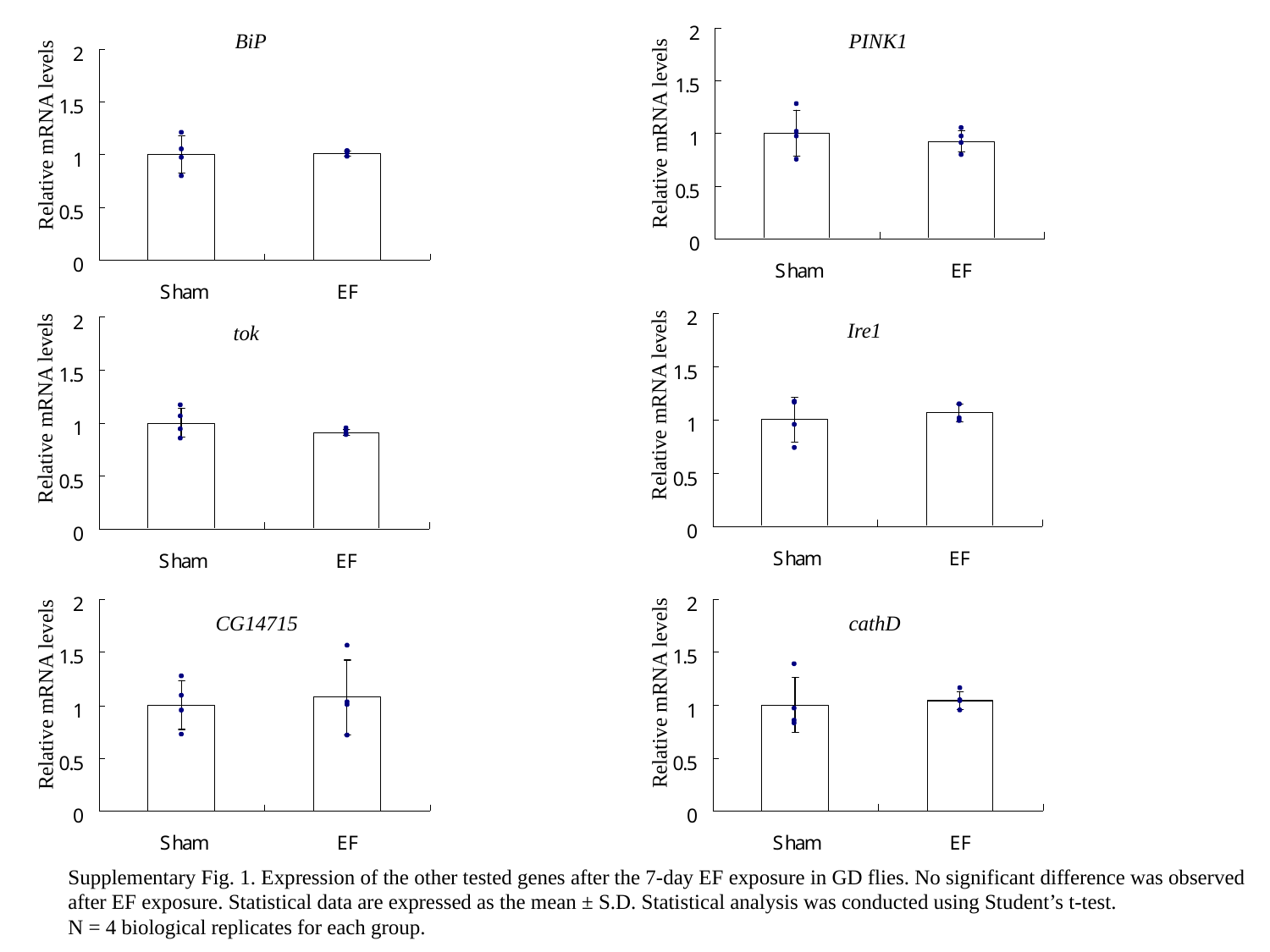

BiP
PINK1
Relative mRNA levels
Relative mRNA levels
Ire1
tok
Relative mRNA levels
Relative mRNA levels
CG14715
cathD
Relative mRNA levels
Relative mRNA levels
Supplementary Fig. 1. Expression of the other tested genes after the 7-day EF exposure in GD flies. No significant difference was observed
after EF exposure. Statistical data are expressed as the mean ± S.D. Statistical analysis was conducted using Student’s t-test.
N = 4 biological replicates for each group.

### Slide 2
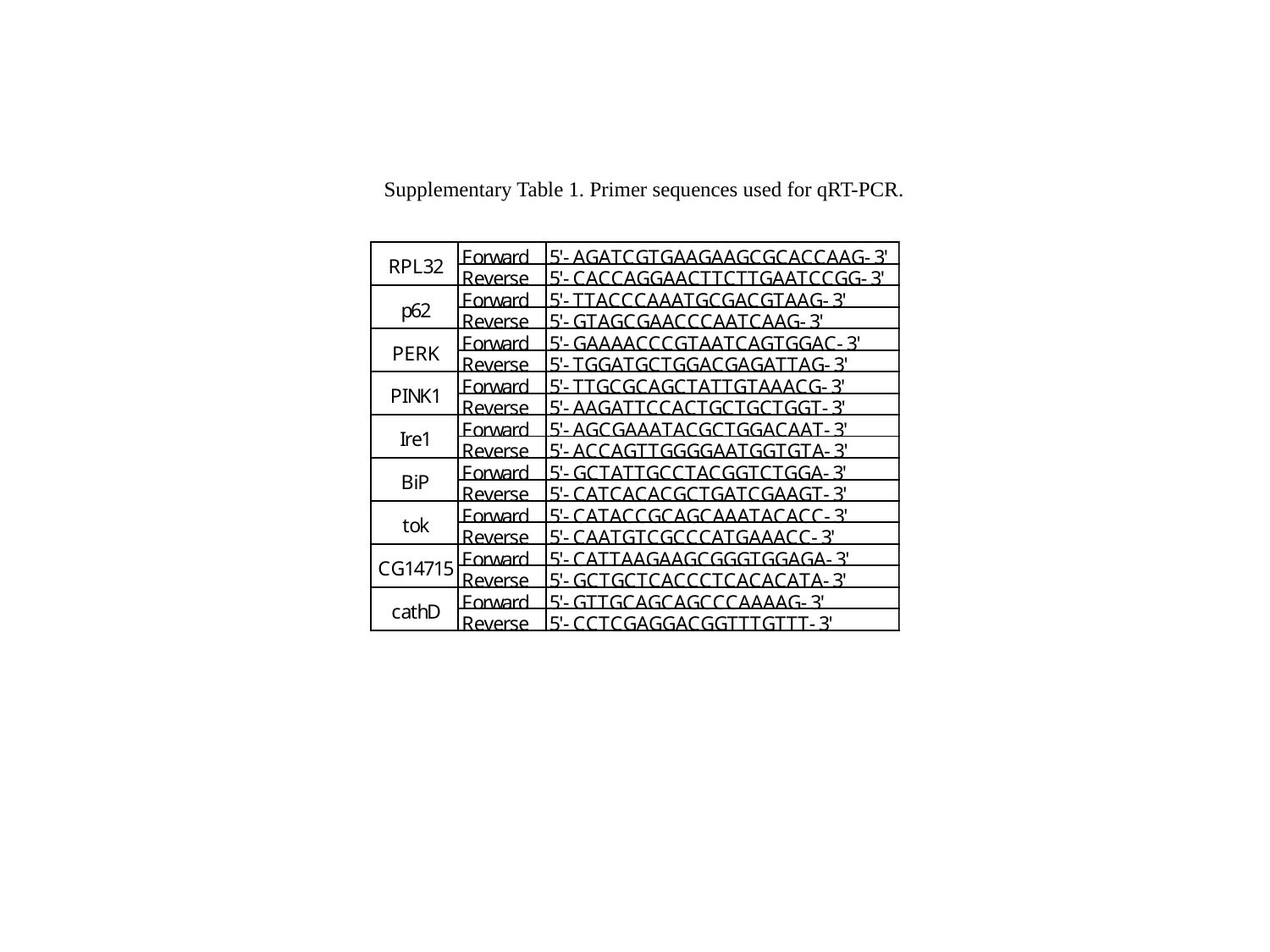

Supplementary Table 1. Primer sequences used for qRT-PCR.
